# Cell-Specific Disruptions in the Basolateral Amygdala-Accumbens Pathway of Fragile X mouse

**DOI:** 10.1101/2023.08.01.551430

**Authors:** Gabriele Giua, Olivier Lassalle, Pascale Chavis, Olivier J. Manzoni

## Abstract

Fragile X syndrome (FXS), a neurodevelopmental disorder caused by the inactivation of the *FMR1* gene, is one of the most common causes of autism spectrum disorder (ASD). People with FXS and ASD often have difficulty regulating their emotions and behaving in a socially appropriate way. The specific neuronal circuits and synaptic dysfunctions involved in social deficits in FXS are not yet fully understood. The basolateral amygdala (BLA) to accumbens core (NAcC) pathway is central to normal social behavior, and we investigated the functionality of this pathway in a novel mouse model of FXS (*Fmr1−/y::Drd1a-tdTomato*). We used optogenetics to distinguish synapses between BLA neurons and spiny projection neurons (SPNs) in the NAcC. We found that the BLA-NAcC connection in FXS mice displayed heightened synaptic strength along with specific alterations in synaptic plasticity. In FXS, enhanced AMPA and NMDA postsynaptic currents elevated BLA-D2 SPN synaptic strength, while BLA-D1 SPNs’ increased firing probability occurred without altered transmission. This results in an overall increase in postsynaptic integration in both D1 and D2 SPN BLA-NAcC pathways compared to WT littermates. Further, the absence of FMRP leads to a specific deficit in long-term depression for BLA-D1 SPNs connections. The neurobiological changes identified in this study may underlie the susceptibility to stressors, socio-communicative difficulties, and altered reward-related processes that are characteristic of FXS pathology.

## Introduction

Fragile X syndrome (FXS) is a prominent neurodevelopmental disorder and a leading monogenic cause of inherited intellectual disability and autism spectrum disorder (ASD)^1, 2^. Affecting around 1.4 out of every 10,000 males and 0.9 out of every 10,000 females^1^, FXS results from the transcriptional silencing of the *FMR1* gene on the X chromosome, causing deficient expression of Fragile X Messenger Ribonucleoprotein 1 (FMRP). This deficiency leads to neurodevelopmental abnormalities in the central nervous system, disrupting neuronal excitability and circuitry and leading to the cognitive and behavioral impairments observed in individuals with FXS^3^. Many aspects of FXS remain poorly understood, particularly regarding the specific brain circuits and synaptic dysfunctions that contribute to the dysregulation of emotional processing and social behavior, which are prominent features of both FXS and ASD.

The nucleus accumbens core (NAcC) acts as a central hub in mesolimbic circuits^4, 5^, playing a pivotal role in integrating emotional, motivational, and reward/aversion signals^6, 7^. GABAergic spiny projection neurons (SPNs) within the NAcC, mainly represented by dopamine receptor 1 (D1R)-expressing SPNs (D1 SPNs) and dopamine receptor 2 (D2R)-expressing SPNs (D2 SPNs), have distinct roles in processing information in healthy conditions and various disorders^8, 9^. The absence of FMRP significantly impacts the intrinsic properties of SPNs and the functionality of their excitatory synapses^10–13^.

Excitatory synapses from the basolateral amygdala (BLA) onto SPNs carry crucial information regarding the emotional valence of stimuli, contributing to a complex network of information processing within the NAcC that guides action selection in reward-related contexts^14, 15^. Activation of the BLA-NAcC pathway has been shown to impair motivation for social interaction, reduce social preference, and induce social avoidance in typical mice^16^. Inhibiting this pathway has been found to restore sociability in a mouse model of autism characterized by social avoidance and impaired social interaction^16^. However, the functionality of the BLA-NAcC connection in the absence of FMRP has not been explored to date.

In this study, the functionality of the connection in the disambiguated BLA-D1 SPN and BLA-D2 SPN synapses was investigated using optogenetics, in vivo electrophysiology, and a novel *Fmr1-/y::Drd1a-tdTomato* mouse model. The results demonstrate synapse-specific alterations in synaptic transmission, postsynaptic integration, and plasticity within the BLA-NAcC connection.

## Methods

### Animals

Animals were treated in compliance with the European Communities Council Directive (86/609/EEC) and the United States National Institutes of Health Guide for the care and use of laboratory animals. The French Ethical committee authorized this project (APAFIS#3279-2015121715284829 v5). Mice were obtained breeding *Drd1a-tdTomato* x *Fmr1* KO2 mice from the Jackson Laboratory (Bar Harbor, ME, USA) and FRAXA foundation respectively. Both strains had a C57Bl/6J background. Mice were acclimated to the animal facility for one week and then housed in male *Drd1a-tdTomato* and female *Fmr1+/-* pairs for breeding. Pups were weaned and ear punched for identification and genotyping at postnatal day (PND) 21. Experiments were performed on first-generation male mice between 70 and 100 PND. *Fmr1+/y :: Drd1a-tdTomato* mice composed the control group (wild type (WT)) and *Fmr1-/y :: Drd1a-tdTomato* the experimental group (knockout (KO)). Mice were housed in groups of 4-5 at constant room temperature (20 ± 1°C) and humidity (60%) and exposed to a light cycle of 12h light/dark with ad libitum access to food and water.

### Stereotaxic surgery

10 μL Hamilton syringe with microinjection needles (32G) was connected to an infusion/withdraw pump (KD Scientific Legato 130) and filled with purified, concentrated adeno-associated virus (2.2×1013 GC/mL) encoding hChR2-EYFP under control of the CaMKIIα promoter (pAAV-CaMKIIa-hChR2(H134R)-EYFP, Addgene 26969-AAV9).

Adolescent male mice (40-50 PND) were deeply anesthetized with isoflurane and treated with carprofen (5mg/Kg, subcutaneous) before placing them in a stereotaxic frame. Microinjection needles were bilaterally placed into the BLA (coordinates from bregma: AP=-1.2mm; ML=±3.1mm; DV=4.8mm) and 250 nL of virus was injected with a speed of 100 nL/min. The needles were left in place for an additional 5 minutes to allow for diffusion of virus particles away from injection site.

### Slice preparation for ex-vivo electrophysiological recordings

Starting from day 30 following stereotaxic surgery, adult male mice were deeply anesthetized with isoflurane and decapitated according to institutional regulations. The brain was sliced (300 μm) on the coronal plane with a vibratome (Integraslice, Campden Instruments) in a sucrose-based solution at 4°C (NaCl 87mM, sucrose 75mM, glucose 25mM, KCl 2.5mM, MgCl2 4mM, CaCl2 0.5mM, NaHCO3 23mM and NaH2PO4 1.25mM). Immediately after cutting, slices containing the NAcC were stored for 1 hour at 32°C in a low calcium artificial cerebrospinal fluid (ACSF; NaCl 130mM, glucose 11mM, KCl 2.5mM, MgCl2 2.4mM, CaCl2 1.2mM, NaHCO3 23mM and NaH2PO4 1.2mM), equilibrated with 95% O2/5% CO2. After 1 hour of recovery, slices were kept at room temperature until the time of recording.

### Electrophysiology

Whole-cell patch clamp of visualized SPNs and extracellular field recordings were made from mouse coronal slices containing the NAcC. Brain slices were placed in the recording chamber and superfused at 2 mL/min with normal calcium ACSF (NaCl 130mM, glucose 11mM, KCl 2.5mM, MgCl2 1.2mM, CaCl2 2.4mM, NaHCO3 23mM and NaH2PO4 1.2mM), equilibrated with 95% O2/5% CO2. The superfusion medium contained gabazine 10μM (SR 95531 hydrobromide; Tocris) to block GABA Type A (GABA-A) receptors. All experiments were done at 25±1°C.

For whole-cell patch clamp experiments, medial NAcC SPNs were visualized using an upright microscope with infrared illumination and then distinguished based on the visualization of *Drd1-tdTomato* using an upright microscope with infrared and fluorescent illumination. Given that in this specific mouse strain, the co-expression of D1 and D2 receptors in SPNs is relatively low (ranging from approximately 1.9% in the dorsal striatum to about 7. 3% in the NAc Core)^17^ it can be reasonably deduced that almost all *Drd1-tdTomato*-positive neurons are D1 SPNs and those *tdTomato*-unlabeled are D2 SPNs. Therefore, for ease of understanding, we use the nomenclature D1 and D2 SPNs along this study recognizing that this classification is tentative.

The intracellular solution was based on K-Gluconate (K-Gluconate 145mM, NaCl 3mM, MgCl_2_ 1mM, EGTA 1mM, CaCl_2_ 0.3mM, Na_2_ATP 2mM, NaGTP 0.5mM, cAMP 0.2mM, buffered with HEPES 10mM), except in the AMPA/NMDA ratio experiments where it was based on CsMeSO_3_ (CsMeSO_3_ 143mM, NaCl 5mM, MgCl_2_ 1mM, EGTA 1mM, CaCl_2_ 0.3mM, HEPES 10mM, Na_2_ATP 2mM, NaGTP 0.3mM and cAMP 0.2mM). Their pH was adjusted to 7.2 and osmolarity to 290-300mOsm. Electrode resistance was 2-4MΩ. Access resistance compensation was not used, and acceptable access resistance was <30MOhms. The potential reference of the amplifier was adjusted to zero prior to breaking into the cell. In V-clamp experiments a −2mV hyperpolarizing pulse was applied before each optically-evoked excitatory postsynaptic currents (oEPSCs) to evaluate the access resistance, and those experiments in which this parameter changed >25% were rejected. oEPSCs were recorded at −70mV except where differentially indicated. Input-Output (I-O) experiments were made by recording the response to increasing optical stimulation intensity (0 to 6mW, steps of 0.5mW). AMPA/NMDA ratios were calculated by measuring evoked oEPSCs (having AMPA and NMDA components) at +40 mV. The AMPA component was isolated by bath application of an NMDA antagonist (APV; 50μM, Tocris). NMDA component was obtained by digital subtraction of the AMPA-oEPSC from the dual component^18^. The oEPSP-Spike coupling (ES-coupling) experiments were performed in I-clamp maintaining the membrane potential at −65 mV. Ten traces were recorded for each step of increasing stimulus intensity (0-10mW, steps of 0.5mW), obtaining mean slope and mean spike probability for each step. oEPSP slopes were measured during the first 2 ms, sorted in 0.5 mV/ms bins, and the firing probability was determined for each bin^19^. For long term depression (LTD) experiments, baseline stimulation frequency was set at 0.1 Hz and plasticity was induced by 10 minutes at 10 Hz stimulation (10’-10Hz)^11^. The intensity of optical stimulation was set at 60 % of that required to achieve maximum response performing an I-O curve (0-6mW, steps of 0.5mW).

For extracellular field recordings, the NAcC was visualized using an upright microscope with infrared illumination. Extracellular oEPSPs were recorded in the medial NAcC with an ACSF-filled electrode. I-O and LTD experiments were conducted as described above. The glutamatergic nature of oEPSPs was systematically confirmed at the end of the experiments using the ionotropic glutamate receptor antagonist CNQX (20μM; NIH), which specifically blocked the synaptic component without altering the non-synaptic component (not shown).

### Optogenetic

Optical stimulation was performed with a 473nm laser (Dragon Laser, Changchun Jilin, China) coupled to a 50μm core glass silica optical fiber (ThorLabs) positioned directly above the slice orientated at 30° approximately 350μm from the recording electrode. At the site of recording discounting scattering a region of approximately 0.05mm^2^ was illuminated that after power attenuation due to adsorption and scattering in the tissue was calculated as approximately 100mW/mm^2^ ^20^. Optically evoked responses were obtained every 10s with pulses of 473 nm wavelength light (0-10mW, 2ms). The stimulating optical fiber was positioned in the dorsal part of the medial NAcC (Fig. 1), dorsomedial to the recording electrode.

**Figure 1.**
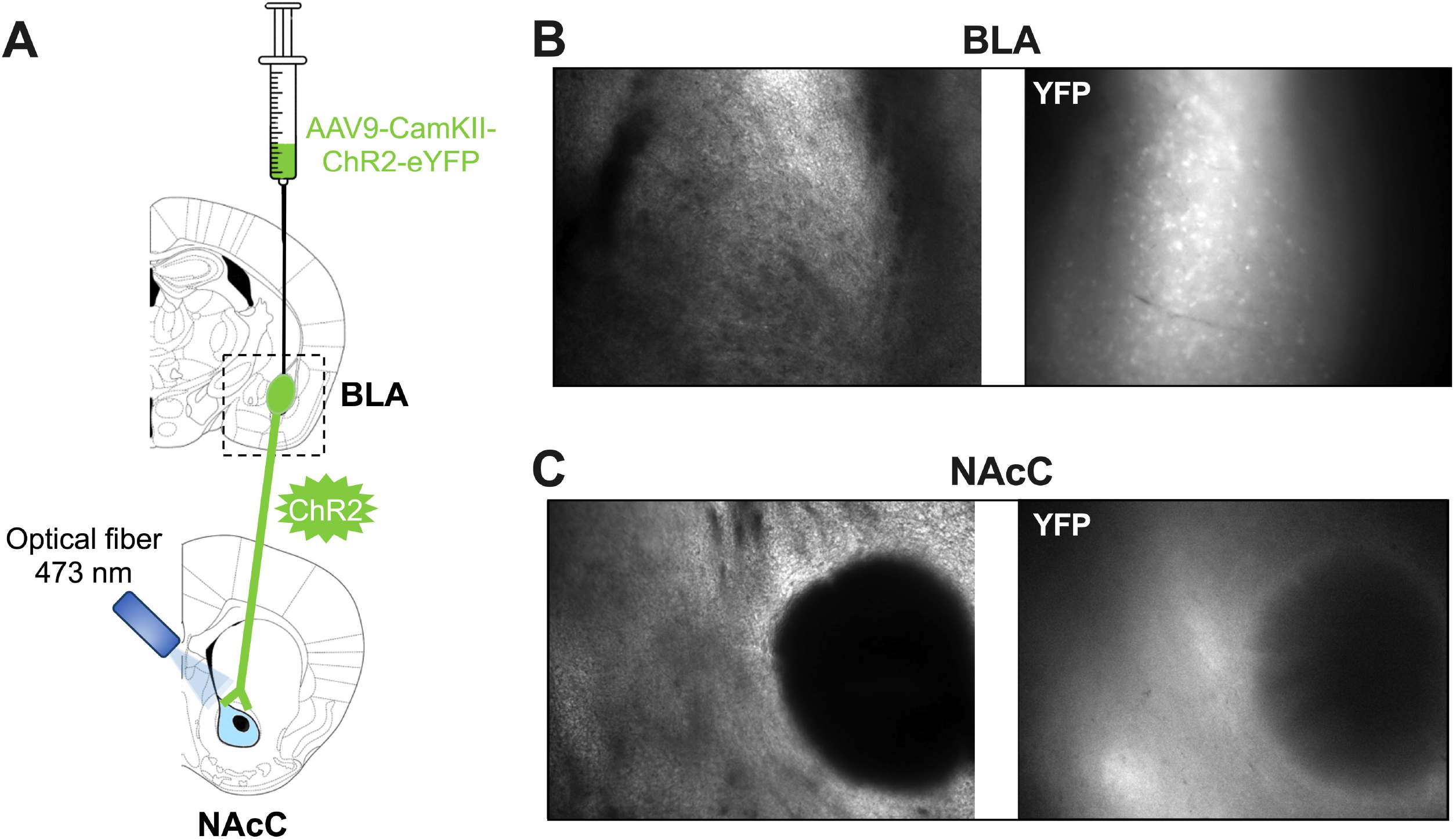
ChR2-eYFP expression in BLA-NAcC pathway. **A:** Schematic illustration of the experimental strategy representing injection of AAV9-CamKII-ChR2-eYFP in BLA, expression of ChR2 along the BLA-NAcC pathway and stimulation of that pathway by optical fiber (473nm). **B:** ChR2-eYFP expression in the injected area (BLA) and **C:** along BLA axons innervating the NAcC (1 month after injection).

### Data acquisition and analysis

All recordings were collected using an Axopatch-200B amplifier (Axon Instruments, Molecular Devices, Sunnyvale, USA), data were low pass filtered at 2kHz, digitized (10kHz, DigiData 1440A, Axon Instruments), acquired using Clampex 10.7 and analyzed using Clampfit 11.2 (Molecular Device).

### Statistics

Statistical analysis of data was performed with Prism (GraphPad Software 9) using tests indicated in the main text after outlier subtraction (ROUT test). N values represent individual animals. Statistical significance was set at *p* < 0.05.

## Results

### FMRP absence selectively increases synaptic transmission from the BLA to the D2 SPNs of the NAcC

Homeostasis of synaptic transmission between the BLA and the NAcC is essential for emotion processing, reward-based learning, addiction, and emotional disorders. The strength of the BLA-NAcC pathway is a key rheostat in autistic symptoms^16^.

We first investigated excitatory synaptic transmission from the BLA to the NAcC in our FXS mouse model by recording extracellular field responses while increasing optical stimulation. The resulting I-O curve revealed a significant alteration in synaptic transmission when FMRP was absent: FXS mice displayed increased synaptic strength in the BLA-NAcC connection compared to WT littermates (Fig. 2). Given FMRP deficiency’s distinct cell subtype specific impact on NAcC’s principal neurons^10^ we next determined if the increased synaptic transmission from the BLA to the NAcC was consistent across D1 and D2 SPNs synapses. Using whole-cell recordings to construct an opto-I-O curve, we found that WT animals exhibited a clear distinction in synaptic strength, with greater transmission from the BLA to D1 SPNs than D2 SPNs (Fig. 3A, E). However, this divergence was absent in KO mice lacking FMRP (Fig. 3B, E). This genotype difference resulted from increased synaptic transmission strength specifically at BLA-D2 SPNs synapses (Fig. 3D, E), with no change observed at BLA-D1 SPNs synapses (Fig. 3C, E).

**Figure 2.**
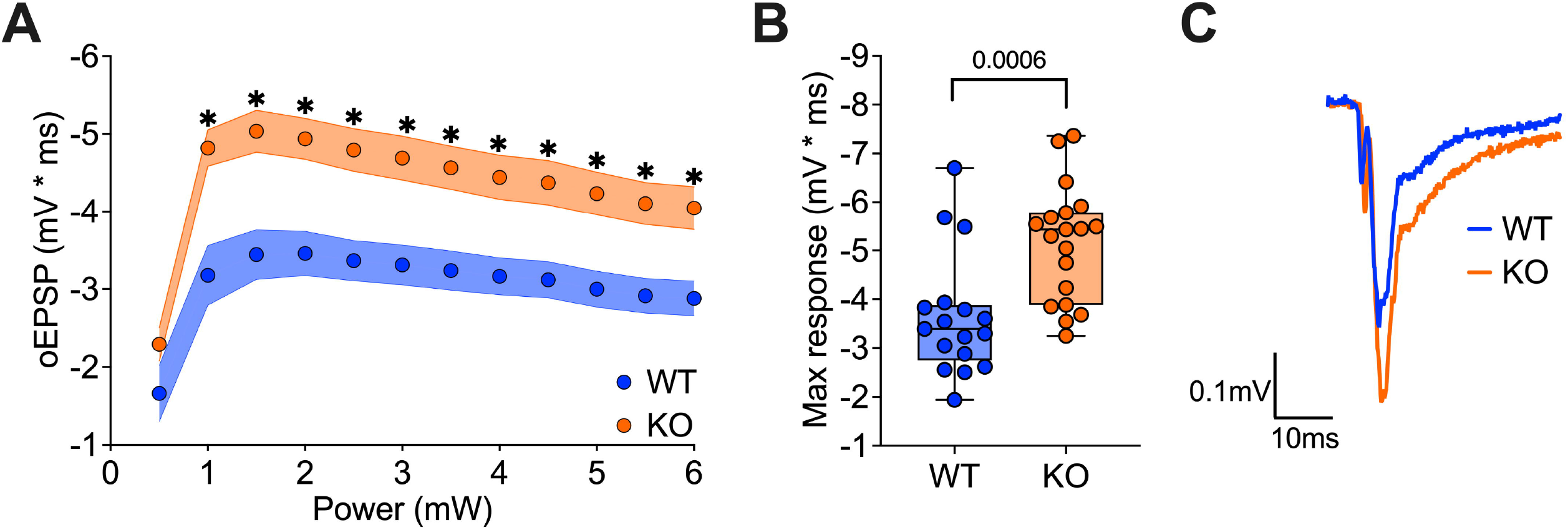
BLA-NAcC synaptic transmission is increased in *Fmr1* KO mice. **A, B:** Extracellular recordings of oEPSPs evoked by increasing light stimulation reveal enhanced synaptic transmission of the BLA-NAcC pathway in *Fmr1* KO compared to WT littermates. **C:** Example trace showing the different oEPSP magnitude elicited by optical stimulation of BLA-NAcC synapses in *Fmr1* WT and KO mice. **A, B:** WT N=17 in blue, KO N=19 in orange. **A:** Single dot represents group mean value at that stimulation intensity step. Data are shown as mean ± SEM in XY plot. Multiple Mann-Whitney *U* test, * p-values <0.05. **B:** Single dot represents an individual mouse. Data are shown as min. to max. box plot with median and 25-75 percentile. Mann-Whitney *U* tests. p-values <0.05 are displayed in graphs.

**Figure 3.**
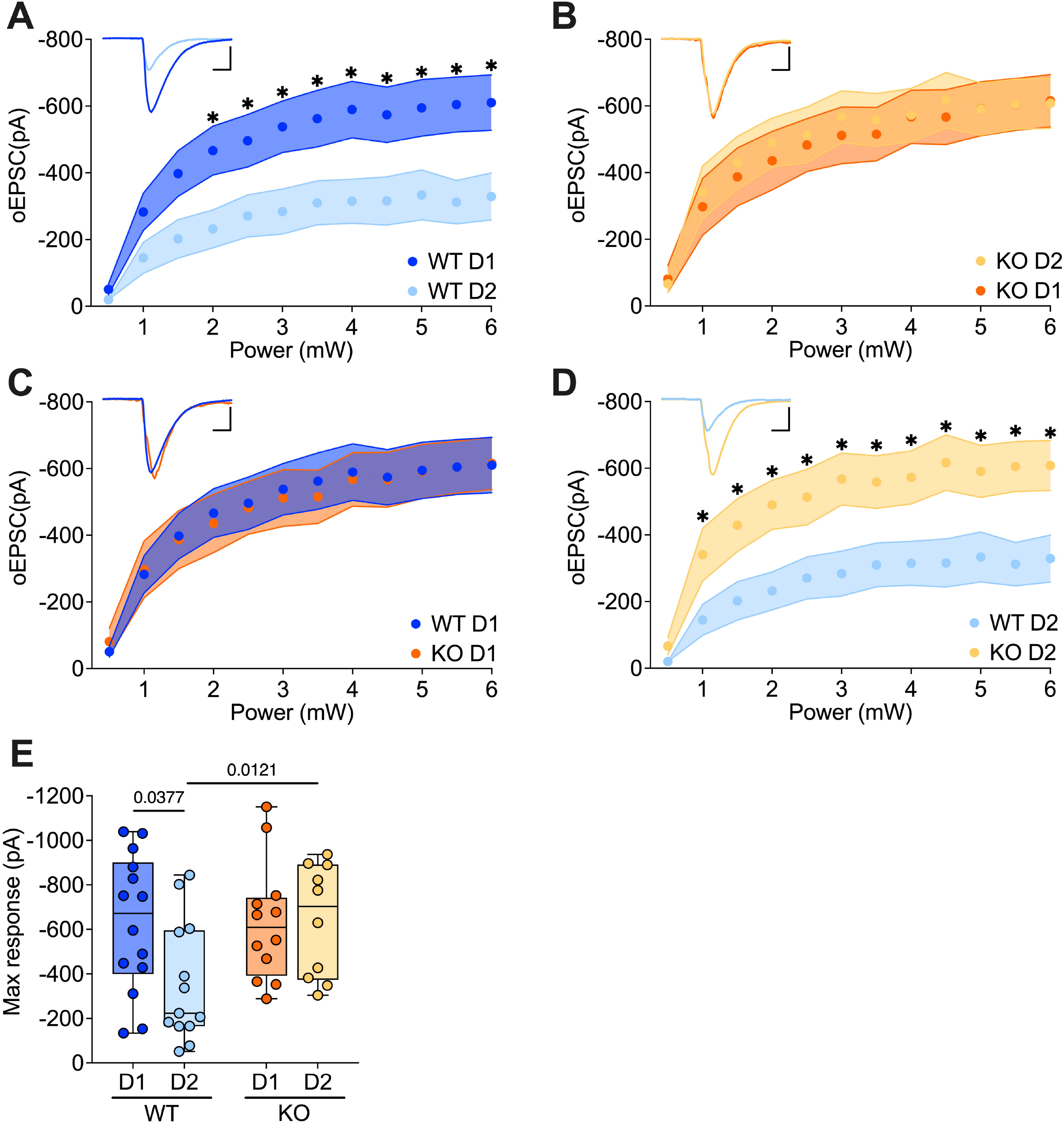
FMRP absence selectively increases the BLA-D2 SPNs pathway. **A, B, E:** Whole-cell recordings reveal that while in *Fmr1* WT the BLA synaptic transmission in NAcC is stronger toward D1 than D2 SPNs, in KO it is equal. **C-E:** BLA-D2 SPNs synaptic transmission is increased in absence of FMRP but not that of BLA-D1 SPNs. **A-E:** WT D1 SPNs N=14 in dark blue, WT D2 SPNs N=13 in light blue, KO D1 SPNs N=11 in dark orange, KO D2 SPNs N=9 in light orange. **A-D:** Single dot represents group mean value at that stimulation intensity step. Data are shown as mean ± SEM in XY plot. Multiple Mann-Whitney *U* test, * p-values <0.05. Example trace in the top left, scale bar: 10ms, 200pA. **E:** Single dot represents an individual mouse. Data are shown as min. to max. box plot with median and 25-75 percentile. Mann-Whitney *U* tests. p-values <0.05 are displayed in graphs.

### FMRP absence equalizes the excitatory receptor repertoire at BLA-D1 SPNs and BLA-D2 SPNs synapses

To investigate potential mechanisms underlying this altered transmission, we evaluated AMPA and NMDA components in both synapse types. The analysis showed that FMRP absence increased both components of BLA-NAcC excitatory transmission exclusively at BLA-D2 SPNs synapses, while BLA-D1 SPNs synapses remained unaffected (Fig. 4B, C). Furthermore, the AMPA/NMDA ratio, a crucial indicator of the balance between synaptic connection strength and potential plasticity, was calculated. In WT animals, a distinct dichotomy emerged: synapses formed by the BLA with D1 SPNs exhibited a lower AMPA/NMDA ratio compared to BLA-D2 SPNs synapses (Fig. 4D). However, this divergence disappeared in the absence of FMRP (Fig. 4D), indicative of a more homogeneous receptor repertoire in KO SPNs.

**Figure 4.**
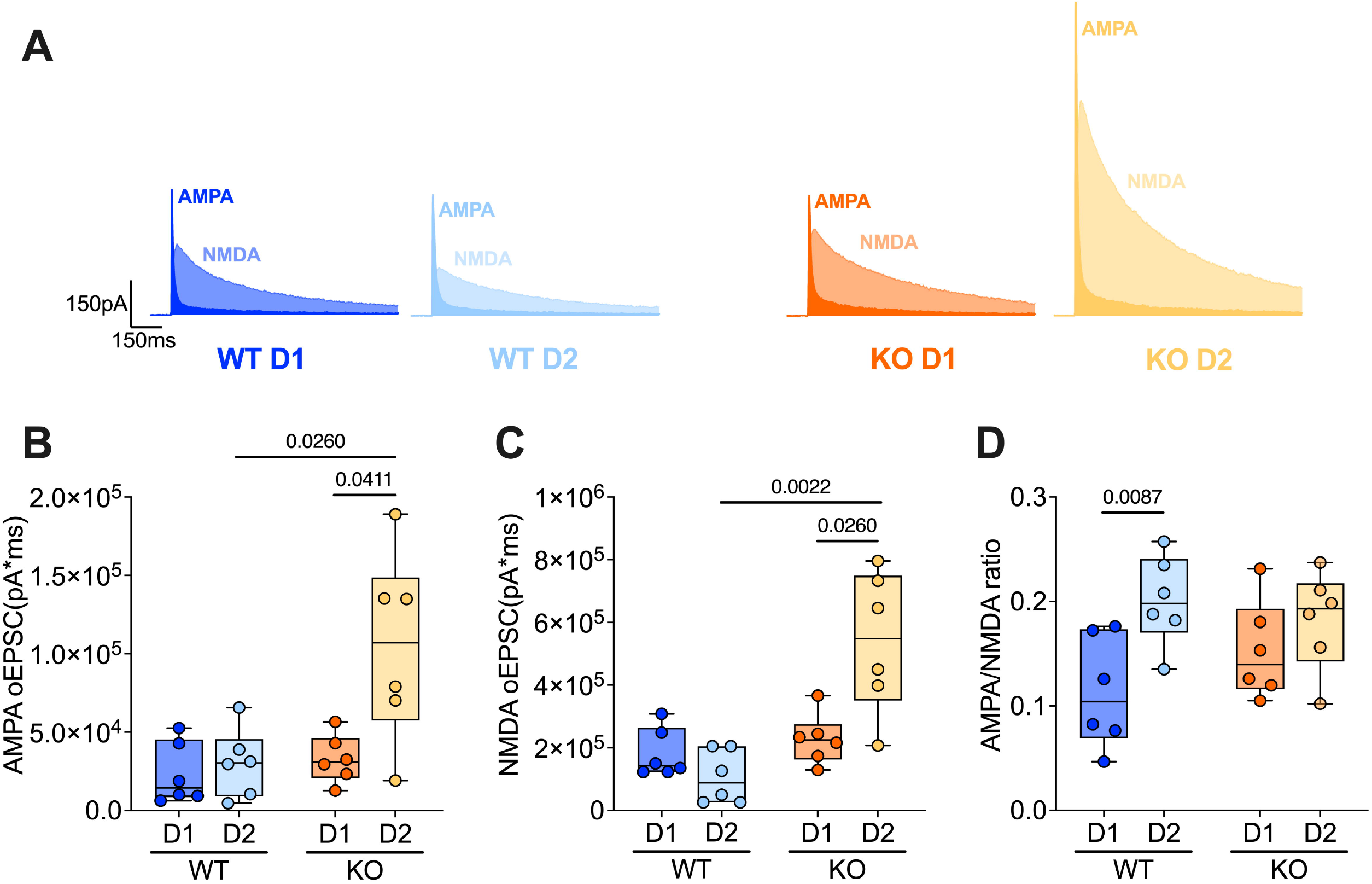
FMRP absence differentially disrupts BLA-D1 and BLA-D2 SPNs postsynaptic receptor repertoire. **A:** Example traces of AMPA and NMDA currents resulting from optical stimulation of BLA-SPNs synapses in *Fmr1* WT and KO littermates. **B, C:** AMPA and NMDA oEPSCs are higher in BLA-D2 SPNs in absence of FMRP. **D:** AMPA/NMDA ratio differs between D1 and D2-SPNs in WTs but not in KOs. **A-D:** WT D1 SPNs N=6 in dark blue, WT D2 SPNs N=6 in light blue, KO D1 SPNs N=6 in dark orange, KO D2 SPNs N=6 in light orange. Single dot represents an individual mouse. Data are shown as min. to max. box plot with median and 25-75 percentile. Mann-Whitney U tests. p-values <0.05 are displayed in graphs.

Overall, these results indicate that FMRP deficiency selectively increases synaptic transmission from the BLA to the D2 SPNs of the NAcC.

### FMRP absence impacts synaptic integration and action potential generation in BLA-NAcC synapses

Synaptic excitatory inputs are integrated within the postsynaptic neuron, orchestrating the initiation of an action potential with a distinct probability contingent upon the synaptic transmission strength and intrinsic neuronal properties, reflecting the synapse’s efficacy. We examined the impact of FMRP’s absence on the integration functionality in synapses between the BLA and NAcC SPNs. Increasing optical stimulation steps were applied to explore the likelihood of action potential generation resulting from the depolarization of SPNs induced by BLA excitatory transmission (Fig. 5A). First, to examine the impact of FMRP absence on the probability of generating an action potential, an ES-coupling graph was built, plotting the probability of firing against the oEPSP slope. This approach allowed the evaluation of whether FMRP influences the probability of action potential generation at the same magnitude of depolarization, thus excluding the influence of different synaptic strengths.

**Figure 5.**
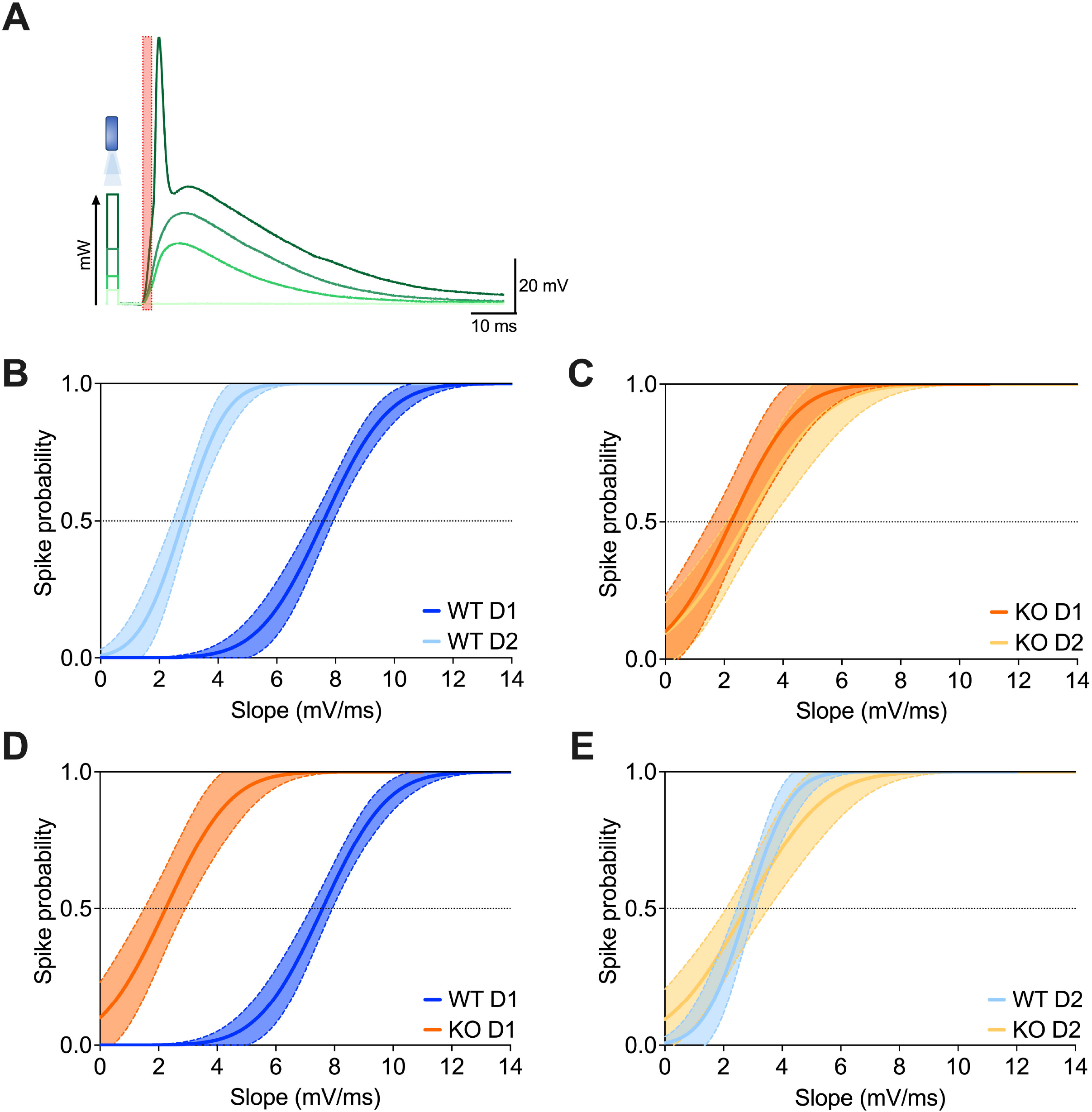
The lack of FMRP increases spike probability in the BLA-D1 SPNs pathway. **A:** Example traces demonstrate that depolarization amplitude increases with optical stimulation intensity until the action potential threshold is reached; the red box indicates the slope extrapolation interval. **B, C:** In *Fmr1* WT mice, stimulating BLA-NAcC synapses, there is a higher probability of triggering an action potential in D2 than in D1 SPNs given the same depolarization magnitude, whereas in KOs this discrepancy is absent. **D, E:** The absence of FMRP induces an increased probability of firing in BLA-D1 SPNs but not in BLA-D2 SPNs synapses. **B-E:** WT D1 SPNs N=6 in dark blue, WT D2 SPNs N=5 in light blue, KO D1 SPNs N=6 in dark orange, KO D2 SPNs N=6 in light orange. Data are presented as non-linear fitting regression curves ± 95% CI in XY plot

Intriguingly, in WT mice, a marked dichotomy emerged, wherein D2 SPNs exhibited a higher half-maximal probability of generating action potentials compared to D1 SPNs at the same oEPSP magnitude (Fig. 5B). However, in FXS mice, both types of synapses demonstrated similar probabilities of inducing action potentials (Fig. 5C). Analyses per cell-type revealed that this discrepancy between the two genotypes stemmed from the fact that the absence of FMRP increased the firing probability in D1 SPNs (Fig. 5D) but not in D2 SPNs (Fig. 5E).

Subsequently, the integration between synaptic strength and firing probability was investigated in BLA-NAcC synapses of *Fmr1* WT and FXS mice. Thus, the firing probability of SPNs was plotted against the intensity of optical stimulation in the BLA-NAcC pathway. The results revealed that D2 SPNs achieved half-maximal probability at lower stimulation intensities compared to D1 SPNs. This distinction was particularly pronounced in WT animals (Fig. 6A) compared to KOs (Fig. 6B). Interestingly, in the absence of FMRP, both D1 and D2 SPNs attained half-maximal probability at lower intensities than their WT counterparts (Fig. 6C, D).

**Figure 6.**
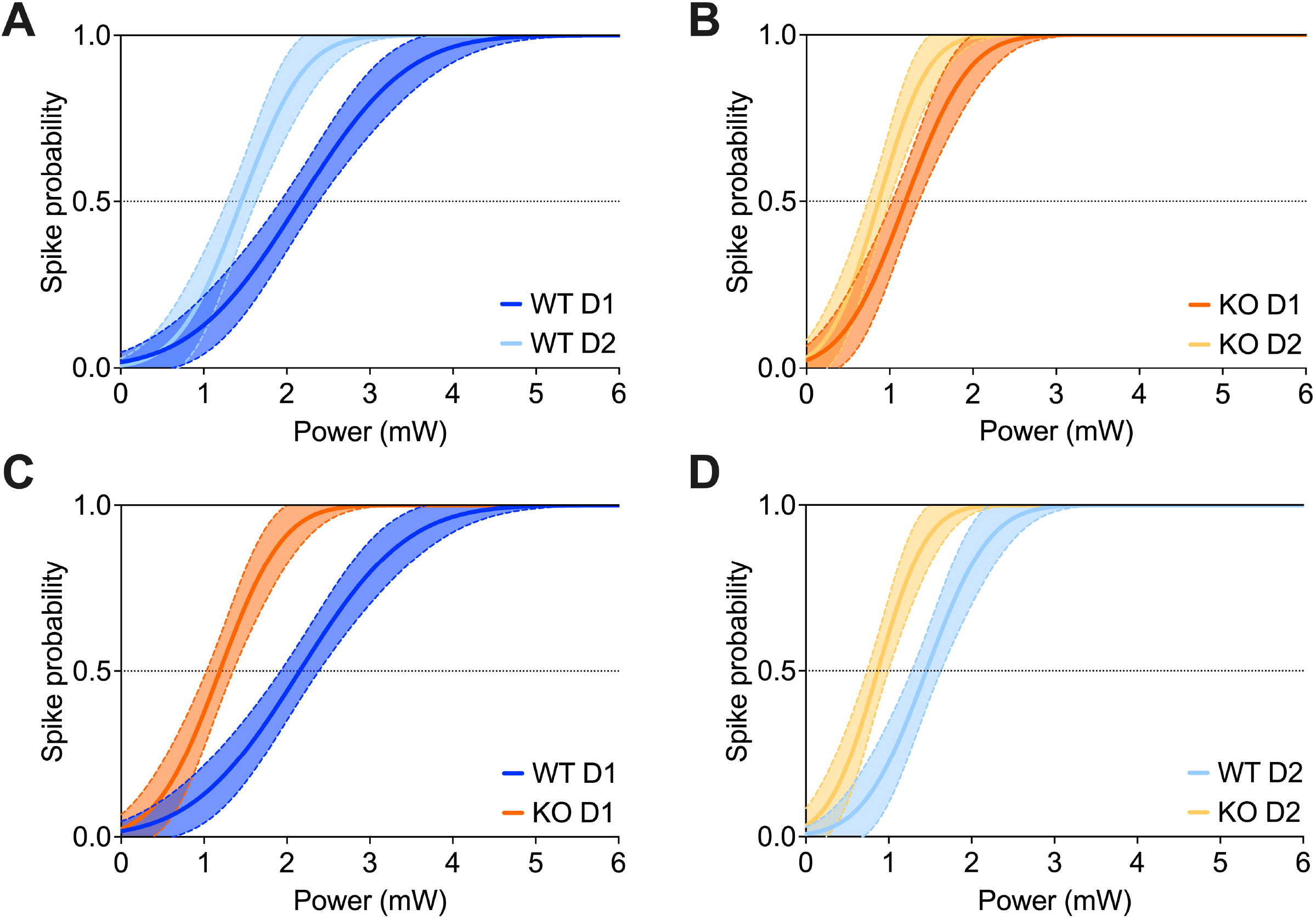
In FXS mice, the spike probability at the BLA-NAcC pathway is increased. **A, B**: D2 SPNs exhibit a higher propensity for firing compared to D1 SPNs under equivalent levels of BLA-NAcC optical stimulation, with this difference more pronounced in WT mice. **C, D:** In the absence of FMRP, both D1 and D2 SPNs show an elevated probability of firing under the same level of optical stimulation from BLA to NAcC. **A-D:** WT D1 SPNs N=6 in dark blue, WT D2 SPNs N=5 in light blue, KO D1 SPNs N=6 in dark orange, KO D2 SPNs N=6 in light orange. Data are presented as non-linear fitting regression curves ± 95% CI in XY plot.

Collectively, these findings demonstrate that in FXS animals, in contrast to WTs, the activation of BLA-NAcC synapses is more likely to result in action potential generation in both D1 and D2 SPNs.

### Selective alteration of LTD at BLA-D1 SPNs synapses in FXS mice

Excitatory synapses in the NAcC of FXS mice have been shown to exhibit impairment in inducing regular endocannabinoid-mediated LTD (eCB-LTD)^11^. The specificity of this deficit to particular inputs remains unknown, but understanding this aspect could provide crucial insights into the FXS phenotype. Consequently, we compared eCB-LTD in BLA-NAcC synapses of FXS and WT littermates. Extracellular field response after 10’-10Hz stimulation revealed that the BLA-NAcC pathway in WT mice exhibits eCB-LTD of synaptic transmission, whereas this ability is impaired in FXS mice (Fig. 7). To investigate the specificity of this phenomenon, disambiguated BLA-SPNs synapses were examined using the whole-cell configuration (Fig. 8). In WT mice, both BLA-D1 SPNs and BLA-D2 SPNs synapses exhibited eCB-LTD (Fig. 8G). However, FMRP-deficient mice showed a deficit in plasticity specifically in the BLA-D1 SPNs synapses (Fig. 8H). This implies that in FXS mice, there is an imbalance of plasticity between BLA-D1 SPNs and BLA-D2 SPNs synapses, differing from what is observed in WTs (Fig. 8A, B, I). These findings collectively confirm that the eCB-LTD deficit observed in the NAcC region in the absence of FMRP also occurs in the specific BLA-NAcC connections. Moreover, this alteration can be specifically attributed to altered plasticity at the BLA-D1 SPNs synapses.

**Figure 7.**
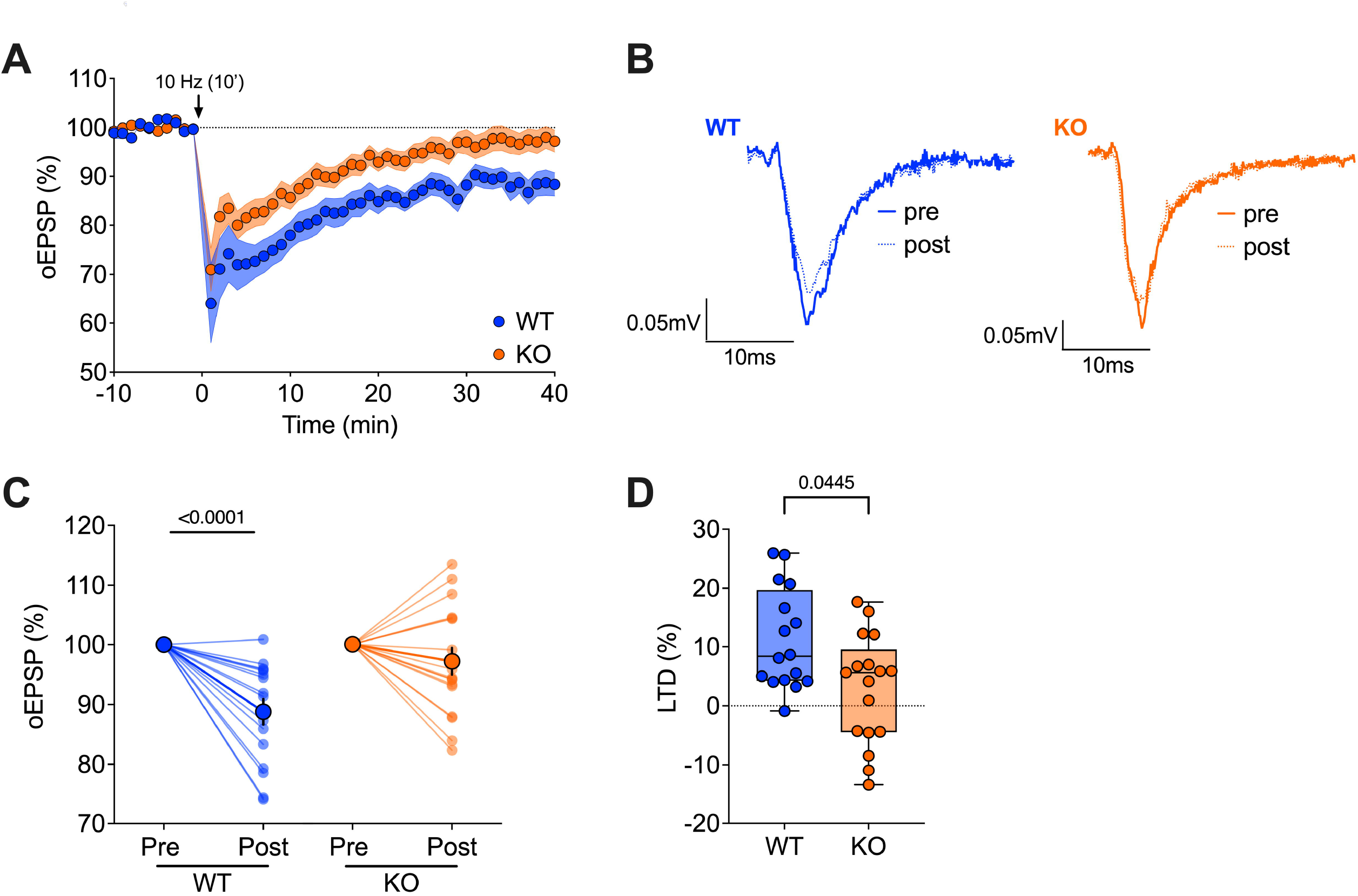
eCB-LTD is completely absent at BLA-NAcC synapses in FXS mice. **A:** Average time-courses of mean oEPSPs illustrate that 10’-10Hz stimulation (indicated by the arrow) induces significant LTD at BLA-NAcC synapses in *Fmr1* WT mice but not in KO mice. **B:** Example traces demonstrate the difference between baseline (solid line) and the response 30-40 minutes after 10’-10Hz stimulation (dashed line). **C:** Pre-post analyses comparing oEPSPs at baseline and 30-40 minutes after 10’-10Hz stimulation reveal the lack of LTD in *Fmr1* KO mice. **D:** The percentage decrease in LTD at BLA-NAcC synapses due to the absence of FMRP. **A, C, D:** WT N=16 in blue, KO N=17 in orange. A: Data are presented as mean ± SEM in an XY plot. **C:** Data are shown as pre-post individual experiments and group average ± SEM, using the Paired Wilcoxon test, with p-values <0.05 displayed in graphs. **D:** Data are shown as a minimum-to-maximum box plot with median and 25-75 percentile, analyzed with Mann-Whitney U tests, and p-values <0.05 are displayed in graphs.

**Figure 8.**
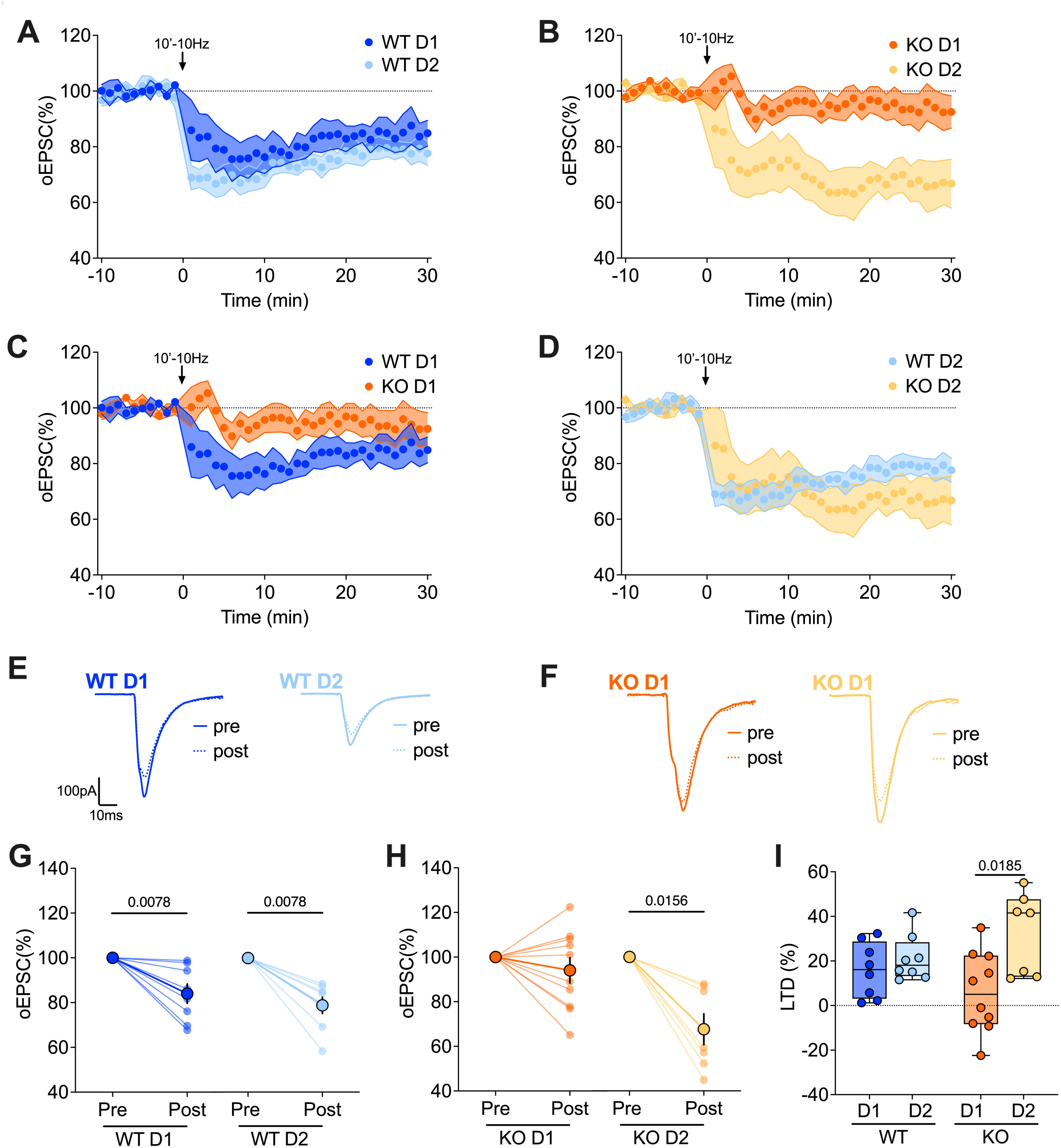
Lack of eCB-LTD in FXS mice is specifically observed in the BLA-D1 SPNs pathway. **A-D:** Average time-courses of mean oEPSCs demonstrate the LTD induced by 10’-10Hz stimulation (indicated by the arrow) at BLA-NAcC synapses per genotype **(A, B)** and cell-type **(C, D)**. **E, F:** Example traces illustrate the difference between baseline (solid line) and the response 20-30 minutes after 10’-10Hz stimulation (dashed line). **G, H**: Pre-post analyses comparing oEPSCs at baseline and 20-30 minutes after 10’-10Hz stimulation reveal that LTD is specifically absent in BLA-D1 SPNs synapses of *Fmr1* KO mice. **I:** While the percentage of LTD remains consistent in the different BLA-SPNs synapses of WT mice, in KO mice, BLA-D2 shows a higher percentage than BLA-D1 synapses. **A-D, G-I**: WT D1 SPNs N=8 in dark blue, WT D2 SPNs N=8 in light blue, KO D1 SPNs N=10 in dark orange, KO D2 SPNs N=7 in light orange. **A-D:** Data are presented as mean ± SEM in an XY plot. **G, H:** Data are shown as pre-post individual experiments and group average ± SEM, using the Paired Wilcoxon test, with p-values <0.05 displayed in graphs. **I:** Data are shown as a minimum-to-maximum box plot with median and 25-75 percentile, analyzed with Mann-Whitney U tests, and p-values <0.05 are displayed in graphs.

## Discussion

We observed that the excitatory glutamatergic transmission between the BLA and NAcC is significantly stronger in FXS mice compared to normal mice. This finding aligns with previous research indicating that in typical animals, activation of this pathway leads to reduced motivation for social interaction, decreased social preference, and increased social avoidance^16^. Conversely, inhibiting this pathway in the Shank3 has been shown to restore social avoidance^16^. Thus, the strength of BLA-NAcC synaptic transmission could serve as a crucial regulator of social approach/avoidance behaviors in both healthy and diseased conditions. Our data indicate that the enhanced glutamatergic transmission observed in FXS mice is specific to BLA-D2 SPNs synapses. Notably, previous research has shown that NAcC D2 SPNs play a significant role in regulating stress susceptibility^21^, and heightened activity in BLA-D2 SPNs is a characteristic feature of mice susceptible to social stress^22^. Therefore, the increased strength of synaptic transmission in BLA-D2 SPNs reported here may contribute to the heightened susceptibility to stressors observed in individuals with FXS.

The glutamatergic BLA neurons that project to the NAcC exhibit a high level of segregation, forming two distinct parallel pathways: one composed of non-CCK neurons projecting onto D1 SPNs and the other comprising CCK neurons projecting onto D2 SPNs^22^. This architectural organization could account for the selectivity we observed in the alterations of synaptic transmission, as the changes might be specific to these two different parallel projections from the BLA to the NAcC. Furthermore, within the same study it was shown that in vivo activation of the CCK^BLA^–D2^NAcC^ pathway has an aversive effect in mice and that social stress selectively activates the CCK^BLA^–D2^NAcC^ circuit^22^. Thus, the increased strength of this specific connection in FXS mice may contribute to the aversive nuances of FXS behavior in stressful or social situations.

BLA hyperexcitability is common in neuropsychiatric and neurodevelopmental disorders characterized by increased vulnerability to stress, enhanced emotional reactivity, or heightened social withdrawal responses^15, 23^. In FXS mice the local GABAergic circuitry of the BLA is altered, leading to hyperexcitability of glutamatergic output neurons^24^. Thus, the increased strength of BLA-NAcC transmission in the absence of FMRP reported here could be due, at least in part, to the hyperexcitability of BLA.

Another potential explanation for altered synapse-specific transmission could lie within the postsynaptic SPNs compartment, which may enable more efficient transmission. FXS models have shown morphological and quantitative alterations of dendritic spines in different brain areas^25^, and previous studies have indicated an increase in the density and length of spines in NAcC principal neurons in the absence of FMRP^12^. Recently, we have shown that the absence of FMRP has a different deleterious effect in the intrinsic properties of D1 and D2, suggesting a cell-specific role of FMRP that could subtend a cell-specific morphological alteration of SPNs^10^. Thus, qualitative and quantitative evaluation of the dendritic arborization and its spines of D1 and D2 SPNs neurons in FXS mice may make an important contribution to the understanding of this phenomenon.

To gain deeper insights into how the excitatory inputs from the BLA are integrated at the postsynaptic level, we conducted ES-coupling experiments. The results revealed that, under the same magnitude of oEPSP, D1 SPNs in FXS mice are more likely to fire compared to those in WT mice, whereas no significant difference was observed in the firing probability of D2 SPNs. This finding is consistent with previous research on the intrinsic properties of SPNs in FXS mice, which demonstrated a lower firing threshold for D1 SPNs in the absence of FMRP^10^, but no such effect was observed for D2 SPNs.

We found that both pathways have a higher likelihood of firing in the absence of FMRP due to two distinct yet converging mechanisms. In the case of BLA-D1 SPNs, while the BLA maintains normal synaptic transmission on D1 SPNs in FXS mice, the absence of FMRP results in a decrease in their firing threshold^10^, thereby facilitating their firing. On the other hand, for BLA-D2 SPNs, although the absence of FMRP does not affect the threshold of D2 SPNs^10^, the BLA exhibits a stronger excitatory synaptic strength, making it more likely for them to reach their threshold and generate action potentials. Indeed, in FXS mice, both D1 and D2 SPNs achieved a half-maximal probability of firing with lower BLA-NAcC stimulations compared to their WT counterparts. These results support the idea that activation of both D1- and D2-mediated pathways in response to BLA stimulation is significantly more likely in FXS mice. This understanding opens avenues for future investigations and potential therapeutic strategies aimed at modulating NAcC to restore the delicate balance disrupted in FXS. By Targeting the dysregulated BLA-NacC circuitry, may help some of the core symptoms of FXS.

Previous research had established that a deficiency of FMRP affects synaptic plasticity in the NAcC^11–13^. A prominent form of plasticity in this brain area is eCB-LTD^26^. While we had previously demonstrated that eCB-LTD is impaired in the NAcC in the absence of FMRP^11^, whether this impairment was pathway specific remained unknown. We found that eCB-LTD was selectively impaired in the BLA-D1 SPNs pathway in FXS. This suggests that FMRP expression is necessary for the proper assembly of the eCB signaling complex at glutamatergic synapses of BLA-D1 SPNs, while it does not appear to be necessary for BLA-D2 SPNs. FMRP is required for the development of cocaine-induced behavioral sensitization, cocaine-related reward learning, and normal NAcC function in these processes^27^. Additionally, selective deficits in eCB-LTD at accumbal D1 SPNs can lead to the suppression of drug and natural reward responses, as well as the seeking of brain stimulation^28^. Together, these findings suggest that the specific alteration of synaptic plasticity in BLA-D1 SPNs within the NAcC of FXS mice may be a biological basis of reward-related behavioral changes in FXS.

Together these findings shed light on a series of intricate dysfunctions within the BLA-NAcC pathway in FXS. Such changes may serve as the biological foundation for specific traits observed in FXS, such as heightened vulnerability to stressors, augmented social withdrawal, and disruptions in reward-related processes.

## Abbreviations

10’-10Hz: 10 minutes at 10 Hz stimulation
ASD: Autism Spectrum Disorder
BLA: Basolateral Amygdala
D1 SPNs: Dopamine receptor 1 (D1R)-expressing SPNs
D2 SPNs: Dopamine receptor 2 (D2R)-expressing SPNs
eCB-LTD: Endocannabinoid mediated-LTD
ES-coupling: oEPSP-Spike coupling
FMRP: Fragile X Messenger Ribonucleoprotein 1
FXS: Fragile X Syndrome
I-O: Input-Output
KO: Knockout
LTD: Long Term Depression
NAcC: Nucleus Accumbens Core
oEPSP: optically-evoked excitatory postsynaptic currents
PND: Postnatal Day
SPNs: Spiny projection neurons
WT: Wild Type

## Declarations

### Ethics statement

Animals were treated in compliance with the European Communities Council Directive (86/609/EEC) and the United States National Institutes of Health Guide for the care and use of laboratory animals. The French Ethical committee authorized this project (APAFIS#3279-2015121715284829 v5).

### Availability of data and materials

All data reported in this paper will be shared by the lead contact upon request. This paper does not report original code. Any additional information required to reanalyze the data reported in this paper is available from the lead contact upon request.

### Authors’ contributions

GG: Conceptualization, Data curation, Formal analysis, Validation, Writing-review and editing.

OL: Data curation, Validation, Methodology.

PC: Conceptualization, Supervision, Methodology, Writing-review and editing

OJM: Conceptualization, Supervision, Funding acquisition, Methodology, Project administration, Writing-original draft, review and editing.

## Acknowledgements

The authors are grateful to the Chavis-Manzoni team members for helpful discussions.

